# A Multimodal Strategy Used By A Large c-di-GMP Network

**DOI:** 10.1101/198481

**Authors:** Kurt M. Dahlstrom, Alan J. Collins, Georgia Doing, Jacyln N Taroni, Timothy J. Gauvin, Casey S. Greene, Deborah A. Hogan, George A. O’Toole

**Affiliations:** Department of Microbiology and Immunology, Geisel School of Medicine at Dartmouth, Hanover, New Hampshire 03755; Department of Systems Pharmacology and Translational Therapeutics, Perelman School of Medicine at the University of Pennsylvania, Philadelphia, Pennsylvania 19104

**Author notes:** These authors contributed equally to this work (Kurt M. Dahlstrom, Alan J. Collins). To whom correspondence should be addressed Department of Microbiology and Immunology, Geisel School of Medicine at Dartmouth Rm 202 Remsen Building, Hanover, NH 03755.

## Abstract

The *Pseudomonas fluorescens* genome encodes for 50+ proteins involved in -di-GMP signaling. Here, we demonstrate that when tested across 188 nutrients, these enzymes and effectors appear capable of impacting biofilm formation. Transcriptional analysis of network members across ∼50 nutrient conditions indicates that altered gene expression can explain a subset, but not all, of biofilm-formation responses to the nutrients. Additional organization of the network is likely achieved through physical interaction, as determined via probing ∼2000 interactions by bacterial two-hybrid assays. Our analysis revealed a multimodal regulatory strategy, using combinations of ligand-mediated signals, protein-protein interaction and/or transcriptional regulation to further fine-tune c-di-GMP-mediated responses. These results create a profile of a large c-di-GMP network that is used to make important cellular decisions, opening the door to future model building and the ability to engineer this complex circuitry in other bacteria.

**Abstract Importance:** Cyclic diguanylate (c-di-GMP) is a key signalling molecule regulating bacterial biofilm formation, and many microbes have up to dozens of proteins that make, break or bind this dinucleotide. Thus, a major open question in the field is how signalling specificity is conferred in this context with a soluble signalling molecule. Here, we take a systems approach, using mutational analysis, transcriptional studies and bacterial two-hybrid analysis to interrogate this network. We find that the network typically combines two or more modes of regulation (i.e., transcriptional control with protein-protein interaction) to generate an observed output.

## Introduction

Cyclic diguanylate (c-di-GMP) is a second messenger used across a wide range of bacterial species to control important life style decisions. c-di-GMP is produced by enzymes called diguanylate cyclases (DGCs) with the canonical GGDEF domain, and is in turn degraded by phosphodiesterases (PDEs) with the canonical EAL domain. This second messenger governs cellular processes by binding to and activating effector proteins that govern such behaviors as biofilm formation, motility, virulence activation, macromolecular synthesis, cell division, and other processes (1-4).

Many bacteria have large c-di-GMP signaling networks, with dozens of DGCs, PDEs,and possible effectors. Several models have been developed to help explain how large c-di-GMP signaling networks may effectively communicate with specific effector proteins at the right time or location in the cell. One model relies on global levels of c-di-GMP signaling to different effectors based on their binding affinity. Pultz *et al.* demonstrated that two *Salmonella Typhimurium* effector proteins with PilZ domains bound c-di-GMP with over a 40-fold difference in affinity (5). This finding was generalized to *Pseudomonas aeruginosa* where over a 140-fold difference was found among its eight PilZ domain-containing proteins (5, 6). Consistent with this “affinity model”, modulating global pools of c-di-GMP has been shown to alter expression and activity of c-di-GMP-metabolizing enzymes, as well as outputs of the network such as exopolysaccharide production and control of flagellar synthesis (7-11). A second model put forth makes use of DGCs, PDEs, and effectors that physically interact with one another allowing for local signaling. Evidence for physical interaction among c-di-GMP enzymes has been found in *Yersinia pestis* (12), and examples of physical interaction being required for signaling include a DGC-PDE interaction in *Escherichia coli* and a DGC-effector interaction in *Pseudomonas fluorescens* (13, 14). Transcriptional control of the genes coding for c-di-GMP related enzymes, including *rapA* in *P. fluorescens* and several DGCs and PDEs in *Vibrio cholerae*, provides a third mechanism of signaling specificity (15-18). Finally, ligand binding may mediate the activity state of various DGCs and PDEs as is the case in *E. coli* for the oxygen-sensing DosC-DosP DGC-PDE pair, or the inactivating effect of zinc on DgcZ (19, 20). Both transcriptional regulation and ligand-mediate activation could conceivable fit in either a global or local signaling model. Thus, there does not appear to be any single mechanism used by bacteria to assure a specific output from this complex network.

In order to understand how large c-di-GMP networks are organized and utilized, we focused on *P. fluorescens* Pf01. The mechanism of biofilm formation in *P. fluorescens* is one of the best understood of such signaling systems, from input to output (21, 22). The effector protein LapD binds cytoplasmic c-di-GMP that in turns leads to an accumulation of the large adhesion LapA on the cell surface via an inside-out signaling mechanism (23, 24). *P. fluorescens* contains a large c-di-GMP network, with 21 GGDEF domain-containing proteins, 5 EAL-domain containing proteins, and 17 dual-domain proteins containing both a GGDEF and EAL domain (**Figure 1**). These dual-domain proteins may function as either DGCs, PDEs, both, or as effectors that can bind c-di-GMP (23, 25-27). *P. fluorescens* also contains 6 proteins with PilZ domains, a domain previously characterized as a c-di-GMP binding effector (28, 29). In a previous study, a mutant library of *P. fluorescens* was assayed for a majority of these genes’ impact on biofilm formation on a single medium containing glycerol and tryptone. Four DGCs and five putative PDEs appeared to impact biofilm formation under this minimal medium condition, leaving most enzymes without an apparent function (30). This conundrum of “extra enzymes” is not unique to *P. fluorescens*. Similar results have been obtained in *V. cholerae*, *P.aeruginosa* and *E. coli* (31-33). This observation leads to two important questions. First, can these other enzymes provide fine control over biofilm formation under non-laboratory conditions? And second, in what ways can such a large network based around a diffusible molecule be organized to provide order to the signaling process?

**Figure 1.**
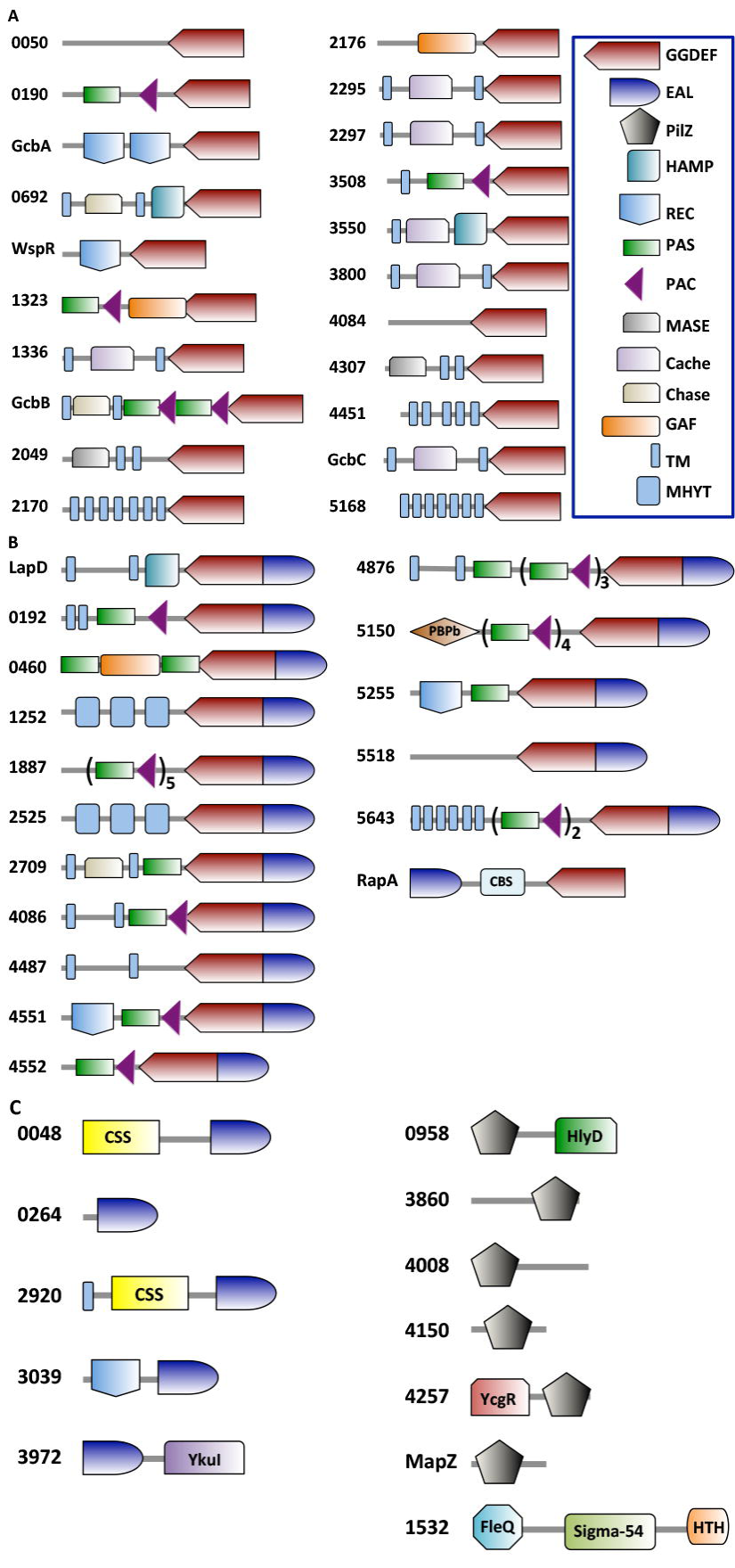
Domain-level depiction of c-di-GMP-related proteins in *P. fluorescens*. All proteins in *P. fluorescens* with a predicted EAL, GGDEF, PilZ, or other known c-di-GMP-binding domain are depicted. Domains illustrated include those predicted by Pfam and SMART. In cases where these two databases predicted different domain architecture over the same amino acid span, the Pfam prediction was used and Pfam domain naming conventions are used in all cases. If a gene/protein name has been assigned/reported, it is also shown here. All recurrent domains are listed in the key, with the remainder of domains labeled on each protein. Models are not drawn to scale, although large amino acid stretches without a domain depicted represent space in the protein where no known domains are predicted. **A.** GGDEF domain-containing proteins. **B.** Dual domain-containing proteins. **C.** EAL, PilZ, and FleQ domain-containing proteins. The domain key is shown boxed.

Here, we attempt resolve these questions using *P. fluorescens* Pf01 as a model system for a large c-di-GMP signaling network. We find that when tested under a broad spectrum of conditions, a large majority of the c-di-GMP-related enzymes and effectors of *P. fluorescens* can impact biofilm formation. Further, we find that transcriptional regulation, non-transcriptional responses to putative ligands and protein-protein interaction are common features among network members. Furthermore, these modes of regulation can be combined, for example, with some pairs of proteins interacting while also responding to common extracellular cues. These findings provide a roadmap for approaching c-di-GMP signaling in bacteria of clinical and environmental relevance, with particular emphasis on understanding how multimodal regulatory strategies modulate biofilm formation in the context of a complex network.

## Results

### A majority of proteins with predicted roles as c-di-GMP metabolizing enzymes or effectors participate in biofilm formation

A previous study from our group constructed a mutant library of a majority of DGCs and PDEs in *P. fluorescens* and tested these mutants for biofilm formation on our standard laboratory minimal medium with glycerol and tryptone (30). Four DGCs – GcbA, GcbB, GcbC, and WspR – were found to positively contribute to biofilm formation under these conditions. Mutations in the genes coding for five dual domain proteins – Pfl01_0192, Pfl01_1887, Pfl01_2709, Pfl01_4086, and Pfl01_4876 – were found to increase biofilm formation under these conditions, indicating that they likely act as PDEs. Given the large array of remaining GGDEF and EAL domain-containing proteins that demonstrated no phenotype under this single growth condition, we hypothesized that some number of these c-di-GMP metabolizing enzymes and/or effectors may also contribute to biofilm formation under conditions not previously tested. Further, in *P. fluorescens* most c-di-GMP related enzymes are predicted to be fused to a variety of ligand-binding domains, including CACHE, CHASE, and GAF domains (**Figure 1**). It therefore seemed possible that the c-di-GMP metabolizing enzymes could be activated by growing *P. fluorescens* under different conditions. To this end, we tested the WT and the 50 strains each carrying a mutation in an individual c-di-GMP-related gene (**Table S1**) using Biolog plates (Biolog, Inc) to test 188 different nutrients (**Table S2**) for their impact on biofilm formation. While we refer to the compounds in each of the wells of the Biolog plates as “nutrients” because they are all organic compounds, these compounds could conceivably serve as carbon sources, input signals, both or neither. The term “nutrients” is used here as a convenient shorthand to generally describe the chemical compounds in the Biolog plates.

The minimal medium typically used to grow *P. fluorescens* biofilms for our studies, referred to as K10-T1 (K10), is a high phosphate medium containing tryptone and glycerol as carbon and energy sources (30). For the purposes of testing biofilm formation in the Biolog plates, we used a similar medium lacking the tryptone with its diverse array of carbon sources. Because the nutrient sources tested here may not be metabolized by *P. fluorescens*, yet may still impact the activity of the c-di-GMP network, glycerol was retained in this medium to permit growth. We refer to this glycerol-containing medium as “base minimal medium” (BMM, see Materials and Methods for details).

In-frame deletions of genes encoding the predicted DGCs, PDEs, and effectors as well as the six genes predicted to encode PilZ domain-containing proteins and a FleQ homologue were constructed. The remainder of tested strains originate from a previously reported (30) single-crossover mutant library (**Table S1**). Each strain was grown overnight in LB with aeration, normalized to OD_600_ of ∼0.716, and added to the BMM with each carbon source found in the Biolog plates before allowing biofilm formation to commence. The nutrients that were toxic to *P. fluorescens*, as indicated by lack of growth of the wild-type strain (not shown), were eliminated from the analysis. Of the remaining nutrients, it was clear that different nutrients have differential effects on biofilm formation (**Figure S1**).

Of the 50 mutants tested, 44 mutants demonstrated a significant difference (p > 0.001) from wild type when comparing their median ability to promote biofilm formation across all nutrients tested (**Figure 2A**). Among strains lacking the GGDEF-containing proteins, 7 showed a significant reduction in biofilm formation as might canonically be expected for this class of mutants. Surprisingly, 11 of these mutants with mutations in DGC-encoding genes showed a significant increase in biofilm formation, suggesting that some putative DGCs may be able to impact biofilm formation contrary to the manner canonically associated with their enzymatic activity. Another 3 mutants in DGC-encoding genes showed no significant impact. Results from strains lacking EAL-containing proteins were more straightforward, with 4 such mutants showing elevated levels of biofilm formation, and a single strain that did not significantly differ from wild type.

**Figure 2.**
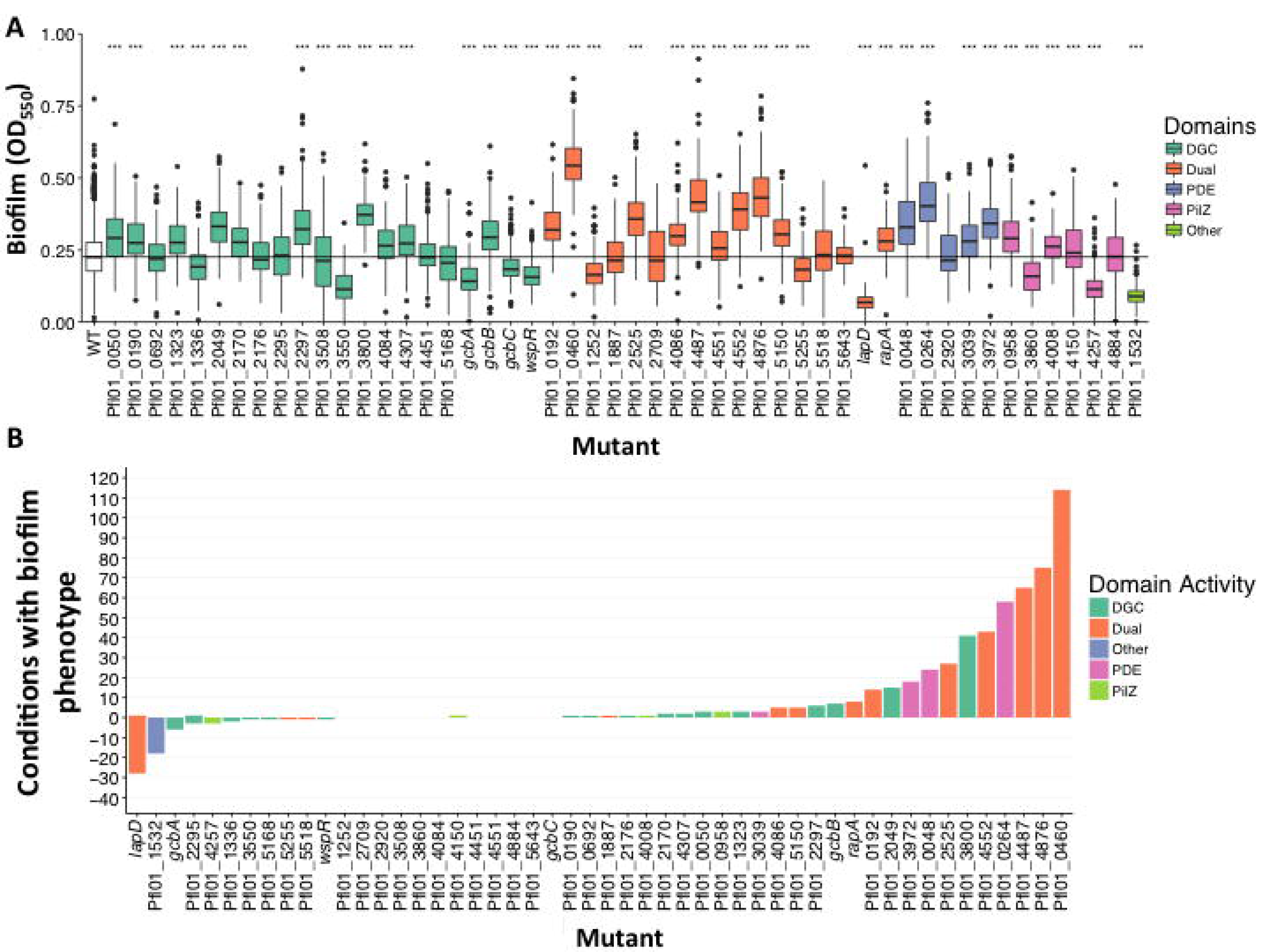
A majority of c-di-GMP-related genes show a defect in biofilm development when mutated. **A**. Data from biofilm formation assays conducted under all 188 tested nutrients for each gene indicated are displayed as a boxplot. Each box is color-coded based on the domain associated with the indicated gene (see legend on right side). Mutants were compared to the wild type using a Wilcoxon signed rank test and p values were adjusted using a Bonferroni correction.***p < 0.001. **B**. Shown is a bar graph of each of the mutants versus the number of nutrient conditions wherein the mutant shows a biofilm phenotype, defined as a value significantly different than the mean WT value using the test described in the Materials and Methods. The height of the bar above and below the origin indicates the number of conditions that each mutant had a significantly enhanced and reduced biofilm, respectively. The bars are color coded to match the domain of the protein encoded by the mutated gene with the legend shown on the right side of the panel.

Analysis of strains lacking the dual-domain proteins revealed a range of results. 10 of the mutant strains demonstrated elevated levels of biofilm formation, three showed a decrease, and four had no significant impact (**Figure 2A**). While prediction of catalytic activity from sequence alone can be unreliable, it is noteworthy that the bulk of mutants lacking dual-domain proteins show elevated biofilm formation (**Figure 2A**). Given that *P. fluorescens* harbors 21 GGDEF containing-proteins while having only five EAL domain-containing proteins, it is perhaps not surprising that most of the dual domain proteins participate in c-di-GMP turnover.

Of the putative effectors, the PilZ domain-containing mutants of Pfl01_0958, Pfl01_4008, and Pfl01_4884 showed significant increases across all carbon sources, mutants of Pfl01_3860 and Pfl01_4257 showed no significant variation, and Pfl01_4150 mutant showed a modest but significant decrease across all carbon sources compared to the wild type. When their encoding genes were mutated, two of the effectors, the FleQ homolog Pfl01_1532 and LapD, both showed significant loss of biofilm formation across a majority of the conditions tested, demonstrating their general relevance to promoting biofilm formation.

Interestingly, 11 mutants that showed no aggregate change in biofilm formation across all conditions. These include the GGDEF domain-containing Pfl01_0692, Pfl01_2176, Pfl01_2295, Pfl01_4451 and Pfl01_5168; the EAL domain-containing Pfl01_2920; the dual domain-containing Pfl01_1887, Pfl01_2709, Pfl01_5518, and Pfl01_5643; and the PilZ domain-containing Pfl01_4884. However, while there may be no global effect detected, there are individual nutrients that appear to impact biofilm formation for these mutants. For example, the Pfl01_1887 mutant showed no significant difference from wild-type in this overall comparison, despite being previously identified as a critical contributor to biofilm formation in this organism in the minimal K10 medium where its mutation results in elevated biofilm formation (30). Taken together, our data suggest that while most genes encoding components of the c-di-GMP network when mutated show a biofilm phenotype under at least some small set of nutrients, other genes appear to play a much broader role in biofilm formation across many environments.

### Biolog data reveals heterogeneity in mutant-by-mutant response to nutrients

To assess trends across nutrient conditions and potential patterns across genes, all values for each genotype were expressed as a ratio relative to that genotype’s median biofilm formation across all conditions. This treatment of the data converts each reading to a measure of how the biofilm in that condition differed from a baseline biofilm formation for that strain. Each of these ratios was then compared to that of WT, and any mutants that show a response to a particular condition as evidenced by a difference in their ratio to baseline by a statistically significant amount (p<0.05, See Materials and Methods for details) are highlighted in **Figure S2** (red: higher ratio than WT; blue: lower ratio than WT; black: not significantly different from WT). We therefore are able to view data points from mutants which differ from how the wild-type behaved in the same carbon source, while downplaying the global effect a mutation might have across all carbon sources from driving the outcome of the analysis (see **Figure 2A**). Squares that are black therefore do not necessarily represent nutrients conditions with no phenotype for that mutant, but represent instances where the mutant did not respond significantly differently from WT to a given condition. This analysis thus provided us an opportunity to examine how mutants behaved in specific nutrients irrespective of their general phenotype.

An immediately apparent finding is the heterogeneity of phenotype that a given mutant can have in particular carbon sources. No class of mutations in GGDEF-, EAL-, Dual- or PilZ- encoding genes was exempt from producing higher and lower biofilm than expected compared to wild type under particular nutrient conditions (**Figure S2**). Further, no class of nutrients tended to generally promote significantly different biofilm levels. The lack of association between particular kinds of nutrients and the phenotypes of particular classes of mutants may indicate that different enzymes in the network respond to different input conditions.

### Differential impact of nutrients as a function of the genes in the c-di-GMP network

To display the relative impact of the different nutrients tested here on biofilm formation by strains carrying mutations in the network, we plotted the percentage of conditions with a biofilm phenotype (decreased or increased) as a function of each mutant (**Figure 2B**). These data show that some mutants display a biofilm phenotype in a large number of the conditions tested. For example, the *lapD* mutant shows significantly reduced biofilm formation in >20% of the conditions tested, confirming its broad importance in biofilm formation. In contrast, other mutants showed changes in biofilm formation in <5% of the conditions tested, with the strain carrying a mutation in Pfl01_5643 showing a biofilm phenotype in only 1 of the 188 nutrients tested. Interestingly, the mutants with increased biofilm formation are not uniformly disrupting genes predicted to code for PDEs, and thus predicted to result in increased c-di-GMP and enhanced biofilm formation. Similarly, the classes of proteins represented on the right side of the figure also include DGCs (**Figure 2B**). These data speak to the differential impact of mutating different components of the network across the possible environments this microbe may encounter.

### c-di-GMP-related genes are transcriptionally regulated in response to nutrients

Given that different DGCs, PDEs and effectors appeared to play a role in biofilm formation depending on the conditions tested, we hypothesized that transcriptional regulation may be partly responsible for determining which enzymes are present to contribute to c-di-GMP signaling. While very few transcriptional studies of DGCs/PDEs/effectors have previously been conducted, there is some precedence for transcriptional regulation. For example, transcription of the *rapA* gene, which codes for a dual domain protein of *P. fluorescens* shown to have PDE activity, has been previously demonstrated to be up-regulated when cells are grown in a low phosphate medium (15).

Our goal was to capture possible roles for transcriptional regulation that may be responsible for phenotypes observed in the Biolog plates. To this end, we ranked carbon sources based on their ability to promote biofilm across all tested strains, and selected 14 “low” and “medium” biofilm-promoting nutrients, 13 “high” biofilm-promoting carbon sources, and 18 “very high” biofilm-promoting carbon sources (**Table S3, Figure S1)**. To assess transcription of all the genes encoding DGCs, PDE, dual domain and PilZ proteins, cells were grown on the BMM with the selected nutrients at a concentration of 2 mM unless, otherwise indicated (see **Table S3**), for six hours before RNA was extracted. Expression analysis was conducted using the Nanostring nCounter system, which directly measures transcript abundance without an amplification step, and raw data were expressed as reads per 1000 counts (see Materials and Methods, **Table S4 & S5**). This normalization strategy has been successfully employed previously in *P. aeruginosa* (34), and we have adopted it here for the purposes of comparing transcripts across many conditions.

To assess the impact of nutrients on expression of the genes in the c-di-GMP network, we plotted the ratio of the transcript reads produced by each genes from each of the 46 nutrients tested over the reads obtained in the BMM (e.g., ratio = reads on nutrient X/reads on base medium). **Figure 3A** shows the individual data points with significant changes in expression as determined indicated by the blue dots (see Materials and Method as for statistical analysis, **Figure S3** for the data summarized as a heatmap, and **Figure S4A** for the figure with a legend showing the nutrients). **Figure 3A** and **Figure S4A** shows that for most genes and for most nutrients, the change in gene expression is <2-fold. However, there are several instances where specific nutrients increase or decrease gene expression from 2-8-fold. We found that the gene showing the highest fold-change encoded RapA, a verified PDE. The expression of *rapA* was previously shown to be strongly up-regulated under low phosphate conditions (15), thus serving to validate our transcriptional analysis. These data indicate that the degree of transcriptional response to any particular nutrient, when such a response is detected, in quite modest (∼2-fold).

**Figure 3.**
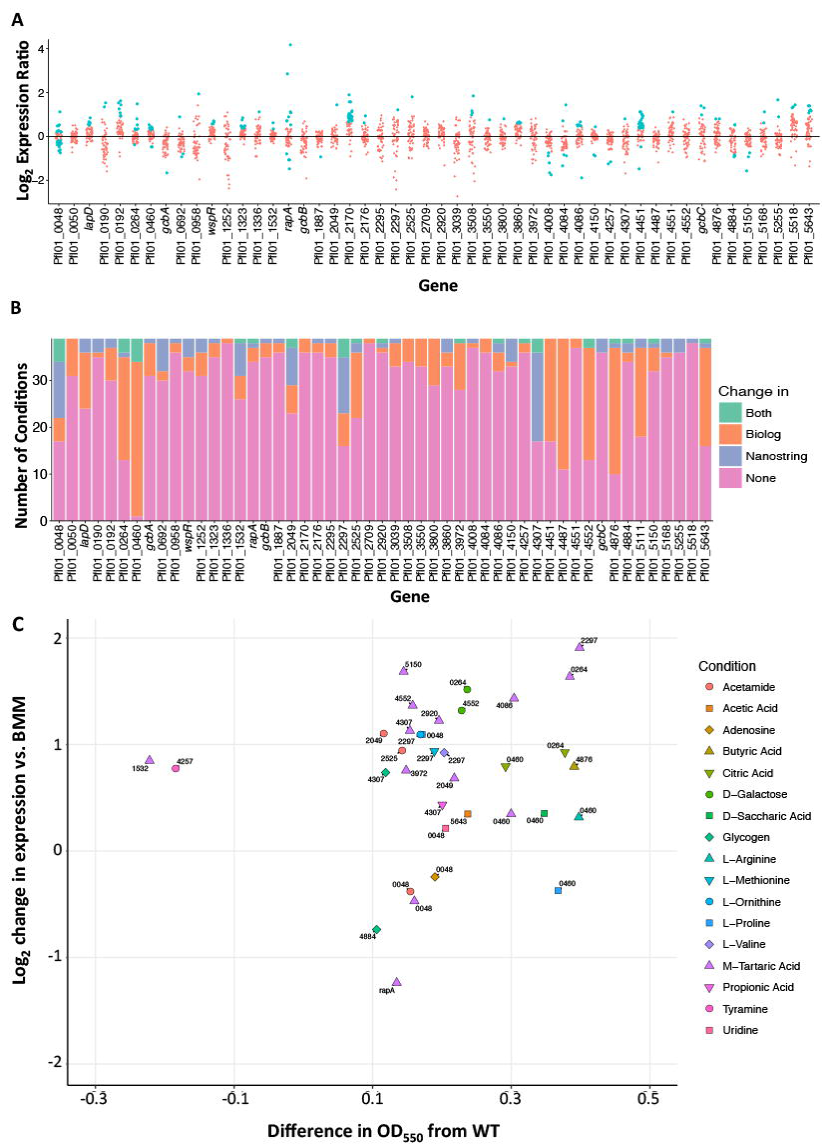
Analysis of the expression of genes in the network. **A**. Shown is the log_2_ expression ratio for each gene in the network across the 50 nutrients tested. The ratio was calculated with dividing the number of counts per 1000 normalized mRNA transcripts on BMM plus the indicated nutrient by the number of mRNA transcripts in the BMM. A positive value indicates an increase in expression in response to the nutrient, while a negative value indicates a decrease in expression. All ratios were transformed to their log_2_ values then plotted. Each red dot indicates a condition in which expression was not significantly different than the expression on BMM alone, while blue dots indicate conditions in which expression was significantly different from BMM (determined as described in the Materials and Methods). A version of this plot with the individual dots assigned to nutrients is shown in **Figure S4A**. **B**. Shown is a plot relating the number of nutrient conditions for which transcription of the indicated gene was changed AND a biofilm defect in a strain carrying a mutation in that same gene was observed in that same nutrient. Data are shown for all the genes in the network. **C.** Conditions where a gene with both a significant change in transcription and the mutation of that gene had a significant impact on biofilm formation are plotted and color coded by nutrient.

### A few nutrients drive the largest changes in expression of genes in the network

In a large network of many related c-di-GMP proteins, one may consider two general methods of gene regulation. The first is to have one or a small number of c-di-GMP-related genes dramatically up- or down-regulated to a given condition, such as the response of *rapA* in low phosphate. An alternative strategy would be to finely regulate a larger number of genes in the network at the same time to produce a desired c-di-GMP output. To test this idea, we examined which nutrients cause the greatest fold-increase and decrease in each gene compared to the BMM. If groups of genes are not regulated together under the same conditions, we would expect that the nutrients causing the largest changes compared to the base medium to be different for each gene.Conversely, if genes are regulated together, we would expect the same carbon sources to be responsible for the largest fold changes in several genes across the network.

We found that six nutrients – m-tartaric acid, glycogen, D-ribose, D-galactose, L-proline, and the low phosphate condition – are responsible for the highest recorded expression of 26 genes in the network (**Table S6** – bolded nutrients). Intriguingly, this same group of nutrients is responsible for the lowest relative expression of an overlapping group of 29 genes. Fold-change compared to the BMM among these genes varied greatly, from near baseline to up to five-fold. Furthermore, m-tartaric acid and glycogen were found to be among the highest and lowest biofilm-promoting nutrients we tested (**Table S3**). Notably, these two nutrients were responsible for a majority of the peak high and low transcriptional values, impacting 18 and 10 genes, respectively (**Table S6**). Overall, 12% of the tested nutrients appear to create the largest transcriptional changes in half of the genes in the network, suggesting that at least for some nutrients that do impact transcription, they may do so across many genes simultaneously.

### Growth on a surface minimally impacts expression of genes in the c-di-GMP network

Previous studies in pseudomonads showed that growth on a surface increases c-di-GMP levels compared to growth as planktonic cells (35-37). Thus, we compared the expression of all genes in the c-di-GMP network for a select set of nutrients in liquid medium versus the same medium solidified with 1.5% agar. Specifically, we investigated gene expression on K10T-1, K10T-1 with low phosphate, BMM, as well as the BMM supplemented with citrate, pyruvate, and L-methionine (**Table S7**). The data were plotted as a ratio of the high c-di-GMP to the low c-di-GMP condition on a per-nutrient basis and analyzed by a Wilcoxon signed rank test with a multiple comparisons correction (**Figure S4B**). Only three genes were found to have a significantly different expression on a surface compared to a liquid (**Figure S4B**). The *gcbA* gene, which has previously been described as being important in early surface attachment (38) was significantly more highly expressed when cells were grown in liquid, while the genes Pfl01_0050 and Pfl01_1336 were significantly more highly expressed when cells were grown on a solid. The *gcbA* and Pfl01_ 1336 genes were both differentially regulated by ∼2-4-fold, while Pfl01_0050 exhibited less than a 2-fold change. Based on the modest dynamic range we observed for most genes in the c-di-GMP network when exposed to different chemical compounds (**Figure 3A**), it may possible that these differences in expression could be sufficient to impact the role of these genes in the response of cells to a surface.

### Modulation of gene expression in the network by c-di-GMP

In *P. aeruginosa*, increased levels of c-di-GMP positively regulate the expression of exopolysaccharide production and down-regulate expression of flagellar genes (8, 10, 39). Furthermore, previous reports have shown c-di-GMP can regulate transcriptional levels of genes through riboswitches, although there have been no reports of such structures in *P. fluorescens* (7, 40). Thus, we assessed the impact of modulating c-di-GMP levels on expression of genes in the network by selecting two strains of *P. fluorescens* that produce low and high levels of c-di-GMP, respectively. The previously reported Δ4DGC mutant lacks the GcbA, GcbB, GcbC, and WspR DGCs, fails to make a biofilm in K10 medium, and produces 2-fold lower c-di-GMP than the wild-type strain (41). Conversely, the GcbC R366E mutant contains a point mutation in the auto-inhibitory site of this DGC, producing over 10-fold more c-di-GMP than the Δ4DGC strain (41). Three biological replicates of each strain were grown on BMM and analyzed via Nanostring (**Table S8**).

Most genes showed changes of <2-fold (**Table S8**), indicating that this large c-di-GMP network does not broadly rely on a core global level of intracellular c-di-GMP to regulate the transcription of other members of the network, although there is a small set of genes whose expression is significant impacted by c-di-GMP levels (**Table S8**, in bold). The gene Pfl01_4451 (the homolog of the *P. aeruginosa sadC* gene) was up-regulated 2.4-fold in the high c-di-GMP mutant compared to wild-type. This result was significant (p<0.01, t-test with Benjamini-Hochberg correction comparing the low and high c-di-GMP producing strains). The *sadC* gene is known to be important in early surface responses. Additionally, Pfl01_0190, Pfl01_5255, and Pfl01_5518 also show significant and greater than 2-fold differences in expression, between the low and high c-di-GMP strains tested here, indicating that transcription of only a small portion of the network is sensitive to large swings in intracellular c-di-GMP concentrations.

### Broad trends observed in the transcriptional network of *P. fluorescens* are also seen in *P. aeruginosa*

Our analysis of the expression of c-di-GMP-related genes (CDG genes) in *P. fluorescens* showed that, with the exception of *rapA* in low phosphate, these genes do not show large variations across the conditions tested. To address whether a similar pattern of expression of CDG genes is found in another related organism, we analyzed the large number of microarray experiments that have been conducted in *P. aeruginosa*. Using the tools developed by Tan *et al.* (42) we assembled a compendium of all published microarrays in *P. aeruginosa*. This compendium includes data for 5549 genes in 1185 experiments including 78 different growth media.

First, to assess whether the broad characteristics of expression of CDG genes in *P. fluorescens* Pf01 are also seen in *P. aeruginosa* we compared the overall expression of CDG genes in both organisms. To do this analysis, we normalised the expression of genes between experiments in the compendium by converting all expression data to values between 0 and 1 according to how they compare to the highest and lowest reported value for each experiment, with the highest value becoming 1 and the lowest value becoming 0. We compared summary statistics of these normalized data to our counts per 1000 normalized Nanostring data (see **Table S5)**. We calculated the median expression for CDG genes in both datasets, as well as the median of non-CDG genes, which we defined as genes coding for proteins without a GGDEF, EAL, HD-GYP, or PilZ domain. We also included the Pho-regulon in the *P. aeruginosa* PAO1 compendium for a comparison with a group of genes that are under robust transcriptional regulation. The pattern of expression observed for *P. fluorescens* Pf01, wherein most genes encoding cdG-metabolizing enzymes show relatively low levels of expression while PilZ-domain-encoding genes are more highly expressed (**Figure S5A**), is also observed for *P. aeruginosa* PAO1 (**Figure S5B**), although the difference in magnitude of expression of the PilZ-encoding genes is lower in *P. aeruginosa* PAO1.

Given that the methods used to assess gene expression differ between *P. aeruginosa* PAO1 (microarrays) and *P. fluorescens* (Nanostring), we sought to use a common metric to compare variation in gene expression between the two microbes. We calculated the coefficient of variance for each data set by dividing the standard deviation by the mean of the expression of each gene across conditions tested. Variability in the expression of these genes across conditions was similar between the two organisms as judges by this metric, with both organisms exhibiting lower variability in CDG gene than the median of non-CDG genes (**Figure S5C,D**). Conversely, the variability of genes in the Pho regulon was higher than that of most non-CDG genes (**Figure S5D**). Together these data suggest transcriptional regulation across multiple growth conditions is not likely a major factor controlling the contribution of genes involved in the c-di-GMP network in either *P. fluorescens* or *P. aeruginosa*.

To investigate potential transcriptional co-regulation common to both organisms, we applied a simple pairwise Spearman correlation analysis of all genes in the compendium and Nanostring datasets. In *P. fluorescens* we found that several genes seem to correlate strongly and significantly with one another (**Figure S5E**), but with fewer genes being significantly and strongly correlated with one another compared to *P. aeruginosa* (**Figure S5F**). These correlation analyses suggest that while the magnitude of transcriptional changes are not large, groups of CDG genes may be under related transcriptional regulation in both organisms.

### Evidence for nutrient-mediated, non-transcriptional control of proteins in the network

We next explored how much of the response of the network to the nutrients tested can be explained by transcriptional regulation. That is, how often does a given mutant’s defect when grown on “nutrient X” correlate with a change in the transcription of that gene when grown on “nutrient X”? To address this question, we assessed what portion of the conditions tested in both Biolog and Nanostring were associated with changes in either biofilm formation, gene expression, both or these phenotypes, or neither (**Figure 3B**).

In most cases that was no link between a change in biofilm formation in a particular mutant and a change in expression of a c-di-GMP-related gene in that condition (**Figure 3B**). However, we did observe instances of biofilm phenotypes that are associated with differences in gene expression (**Figure 3B**), suggesting that gene expression may explain the impact of these conditions. Data from the nutrients with both biofilm impact and gene expression difference are displayed in **Figure 3C**. Interestingly, m-tartaric acid impacted both the biofilm formation and expression of 13 different genes. m-Tartaric acid also induces one of the highest overall biofilm tested, suggesting that m-tartaric acid may be an important signal or nutrient for *P. fluorescens* and exert at least some degree of its impact on biofilm formation via transcriptional control. As noted above, in a majority of circumstances, when biofilm formation occurs it can be attributed to the chemical compound added in the absence of any detectable transcriptional change (**Figure 3B**). Given the large number of putative sensory domains present in many of the c-di-GMP-related proteins in *P. fluorescens* Pf0-1, we suggest that nutrients may be impacting the c-di-GMP network by controlling enzyme activity.

### Physical interaction is common for DGC and dual-domain proteins

While patterns of biofilm formation and transcriptional regulation emerged from the differing nutrient conditions tested, these findings did not appear sufficient to explain how this larger network might be ordered to prevent cross talk among its many enzymes and effectors. Furthermore, physical interaction as a mechanism of signaling specificity has precedence in a number of bacterial c-di-GMP systems including *E. coli*, *P. aeruginosa*, and *Xanthomonas axonopodis*. (13, 43, 44). In *P. fluorescens*, physical interaction between the DGC GcbC and the effector LapD has previously been shown to be required for specificity of GcbC signaling in biofilm formation (14). However, it is largely unknown how common physical interaction is among enzymes and effectors in the network, whether a given protein may have a single or many interaction partners, or if a given type of c-di-GMP protein is more or less likely to form physical interactions than others.

To address these questions, each c-di-GMP related gene in *P. fluorescens* was cloned into both “bait” and “prey” vectors of a bacterial two-hybrid (B2H) system to assess the capacity of each protein for physical interaction. Each member of every protein type – GGDEF, EAL, dual-domain, and PilZ – was tested against all members of the other types. Additionally, all dual-domain proteins were also tested against all other dual-domain proteins as it is possible that such proteins may act as DGCs, PDEs, both or as effectors. The summary of this analysis, which represented close to 2000 bacterial two-hybrid assays, is presented in **Figure 4, Figure S6** and **Table S9**.

**Figure 4.**
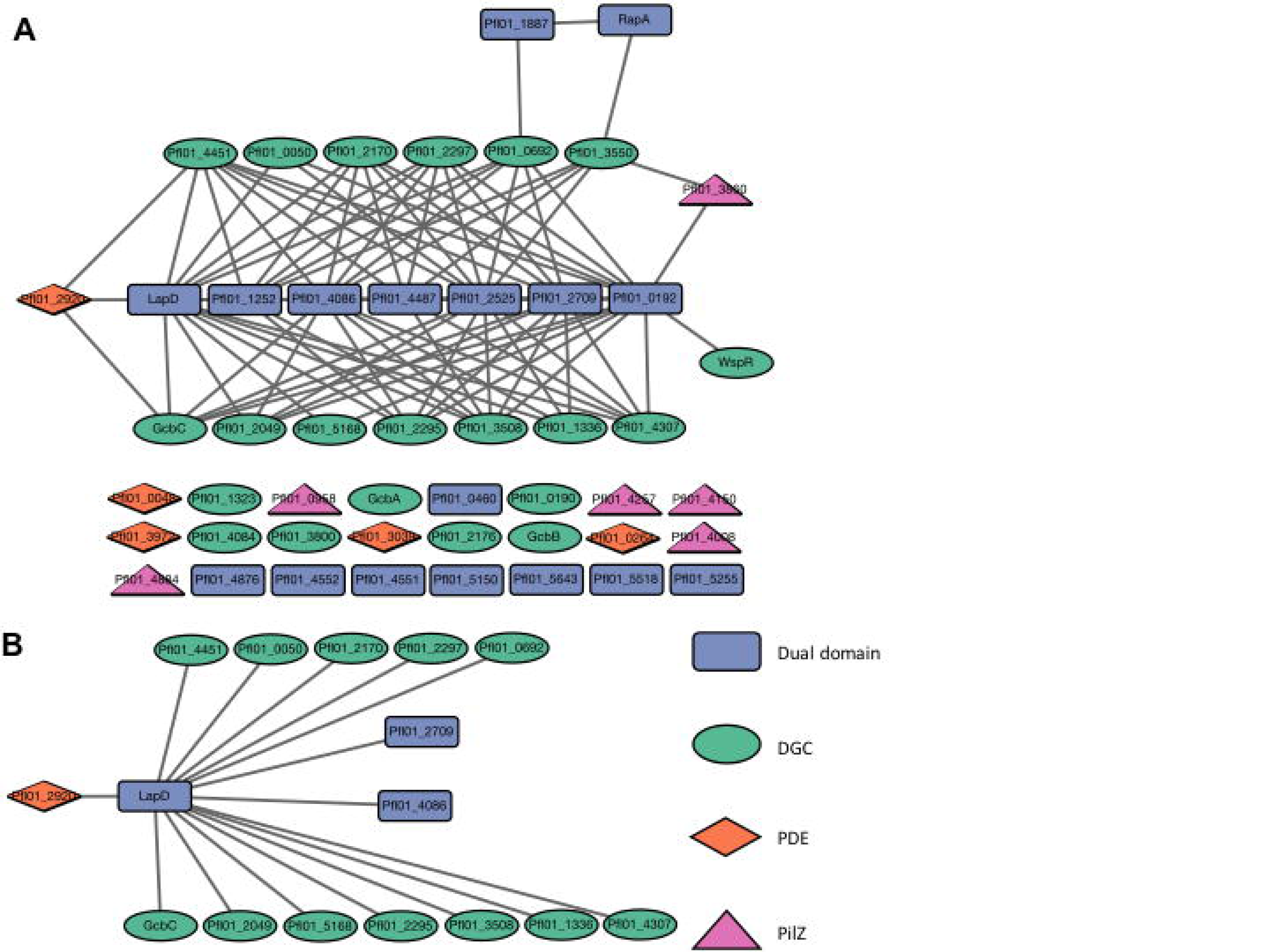
Physical interaction map of c-di-GMP-related proteins in *P. fluorescens*.A. BTH101 *E. coli* cells were grown on LB medium supplemented with X-gal for 20 hours and colonies were assessed for blue color. Protein pairs of each domain type were tested for physical interaction in the bacterial two-hybrid system against each member of every other domain type, with dual-domain proteins also being tested against each other. Each gene was tested on both the pUT18 and pKNT25 vectors; a positive result was recorded if either the T18- or T25-fused protein produced blue colonies with another member of the network. Proteins that failed to interact are shown at bottom. FleQ was not tested in this analysis. The interaction map was generated by Cytoscape v 3.5.1. **B**. LapD serves as a node of interaction among several proteins shown here. The legend is shown on the bottom, right of the figure: GGDEF-containing proteins are represented by green circles, EAL-containing proteins by orange diamonds, dual-domain proteins by blue rectangles, and PilZ-containing proteins by purple triangles.

Several dual-domain proteins showed a relatively high capacity for interaction with other network members (**Figure 4A**). In particular LapD, Pfl01_0192, Pfl01_1252, Pfl01_2525, Pfl01_2709, Pfl01_4086, and Pfl01_4487 interacted with 7 to 18 partners each. A subset of 15 proteins interacted with LapD (**Figure 4B**) and four additional dual-domain proteins interacted with greater than 10 partners (**Table S9 and Figure S6**). Conversely, RapA and Pfl01_1887 are not a part of these larger hubs, interacting with only two and one partners, respectively (**Figure 4A and Table S9**).

DGCs showed a highly variable propensity to interact (**Table S9**). 14 DGCs appear to participate in physical interaction, having between one and eight interaction partners. Interestingly, the four DGCs previously found to be necessary for biofilm formation under K10T-1 media conditions have different interaction profiles. GcbA and GcbB failed to interact with other proteins in this assay, while by contrast GcbC interacts with several dual-domain proteins (including LapD) and with an EAL domain-containing protein. WspR has a single interaction partner - Pfl01_1092. The remaining seven DGCs do not engage in detectable physical interaction under our assay conditions.

Finally, little evidence was found to indicate that EAL and PilZ domain-containing proteins interact with other members of the network (**Figure 4A**). Among the five EAL domain-containing proteins, only Pfl01_2920 was found to interact with two DGCs and LapD. Likewise, a single PilZ domain-containing protein, Pfl01_3860, interacts with the GGDEF domain-containing protein Pfl01_3550 and the dual-domain protein Pfl01_0192. The Pfl01_3860 PilZ domain-containing protein is part of a small node made up of a GGDEF domain and two dual-domain proteins (**Fig. 4A**). Together, these data indicate that a number of the proteins in the network have the potential to interact, and thereby potentially form signaling complexes.

### Relationship between interacting proteins and shared biofilm phenotypes

Having shown that some, but not all impacts of different conditions on biofilm formation may be explained by transcriptional regulation, we next tried to determine how much of a role physical interaction might play in contributing to signaling specificity in the network. To perform this analysis, we assessed the conditions wherein both members of an interacting pair of proteins also displayed a significant biofilm phenotype when mutated. We scored these biofilm phenotypes based on whether they were in the same direction (i.e. both enhanced biofilm or both reduced biofilm) and calculated the proportion of the total biofilm phenotypes each member of the interacting pairs shared (**Figure 5A**). This analysis shows that most interaction partners impact biofilm in only one or two of the same conditions, however some pairs do demonstrate a large overlap. For example, the interaction pairs such as *wspR-*Pfl01_0192 and Pfl01_0692-Pfl01_1887, which only impact biofilm formation in a few conditions, overlap in most of those conditions. Furthermore, there are several instances wherein interacting pairs, when mutated, show opposite biofilm phenotypes in a particular nutrient condition. Such a finding is likely physiologically relevant given that the biofilm formation is impacted under a common growth condition. It is also not surprising that overlapping conditions between interaction partners are generally consistent in being all in the same direction or all in the opposite directions, as mutants generally only impacted biofilm formation in one direction. However, it is worth noting that interaction partners seem equally likely to act on biofilm formation in the same direction as they do in opposite directions. These data suggest that there may be numerous examples of both antagonistic and cooperative interactions in this network, often involving the same proteins, and possibly modulated via physical interaction.

**Figure 5.**
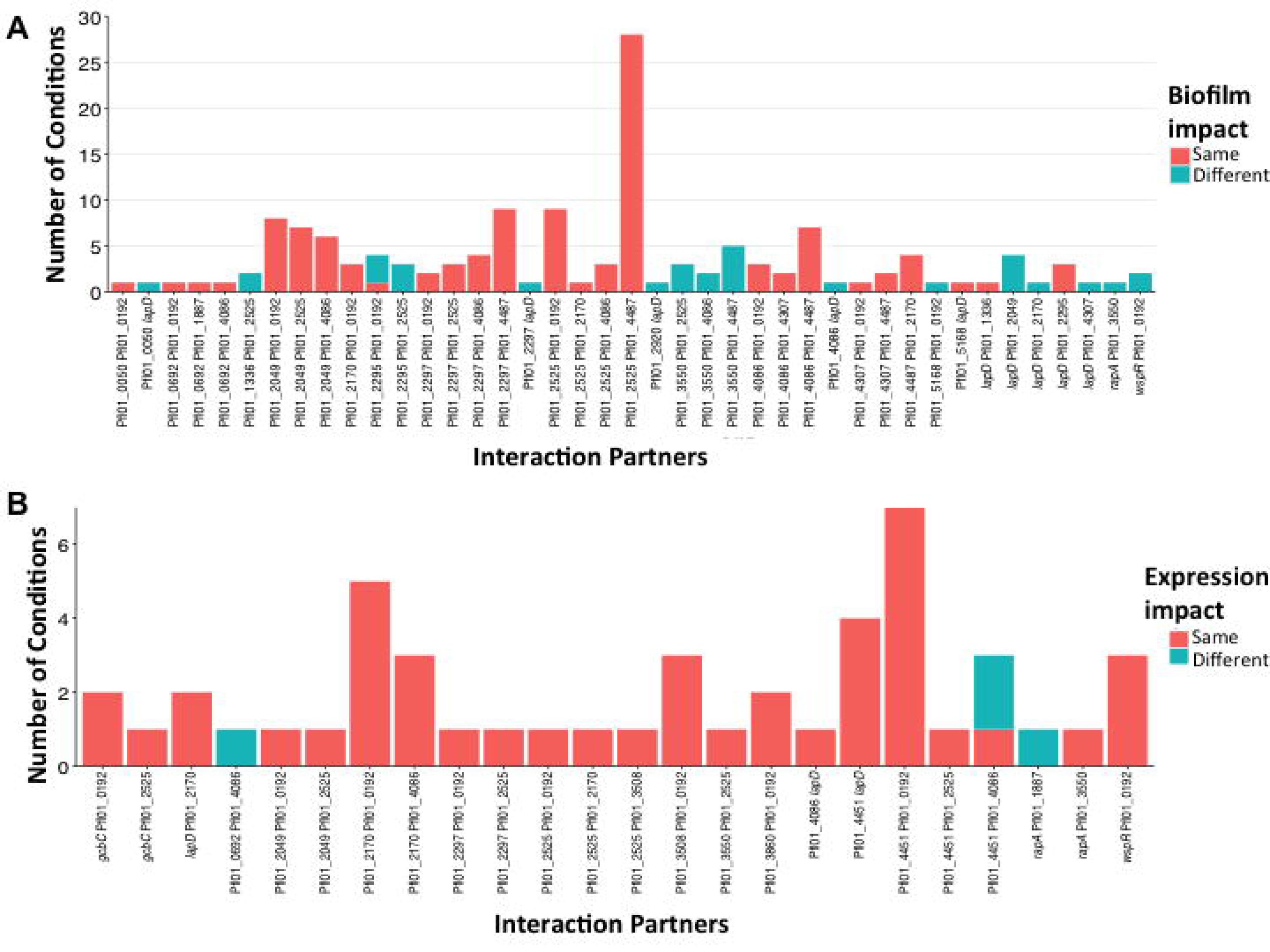
Assessing the protein-protein interactions in the network. **A**. Shown is a plot of interacting pairs of proteins in the network versus the number of nutrient conditions wherein each pair shared a biofilm phenotype. Orange coloration indicates the number of conditions where both interactors affected biofilm formation in the same way, while blue color indicated the number of conditions that interactors affected biofilm formation in opposite directions. **B.** The stacked box plot shows the relationship between interacting pairs of proteins in the network versus a change in expression of both genes in the interacting pair in the same nutrient. Orange coloration indicates the number of conditions where both interactors showed a change in transcription in the same direction, while blue color indicated the number of conditions that interactors showed a change in transcription in the opposite direction. Note the difference in the scale bar between panels A and B.

### Intersection between B2H and transcriptional control

We next examined possible overlap between proteins that interacted with one another and were coordinately expressed. We scored the number of conditions where genes of an interacting pair changed in expression under the same condition. A number of interacting pairs showed coordinated transcriptional changes under one to seven conditions (**Figure 5B, Table S10**). Interesting, some genes that only showed significantly different expression from the BMM in one or a small number of conditions (see **Table S5**) did so under the same nutrient additions as genes encoding for their interaction partner. For example, for the Pfl01_0192-Pfl01_4451 interacting pair, Pfl01_0192 shares all seven conditions that produce differential expression from the BMM with the GGDEF domain-containing Pfl01_4451. Together, these results suggest that in some cases, a combination of transcriptional control and physical interactions may be utilized to modulate signalling specificity.

## Discussion

In this study, we investigated the c-di-GMP network of *P. fluorescens*. We selected this model organism because it has one of the best-understood c-di-GMP circuits, from input to output, with the control of localization of the cell-surface biofilm adhesion LapA as a key output (21, 22). LapA appears to contribute to or be required for biofilm formation under every condition we have examined to date, including the base minimal medium used in this study, and this same medium supplemented with the 188 nutrients investigated here. Our analysis revealed one simple, central conclusion: no one mode of regulation outlined above (nutrient input, transcriptional regulation, protein-protein interaction) adequately describes the function of the network. Instead, these modes of regulation can function in combination to contribute to tuning of the network. We propose that this multimodal strategy of controlling c-di-GMP-mediated outputs ultimately provides fidelity and specificity to the network.

We found that a majority of c-di-GMP related genes in *P. fluorescens* impact biofilm formation when mutated and tested under a large array of conditions, indicating that context is key for understanding the function of these large signaling networks. Further, this context appears to shape the organization of the network in several ways. We found that transcriptional regulation in *P. fluorescens* rarely results in over 5-fold changes in transcription across all tested conditions. It was also rare for any given nutrient and any single gene that a transcriptional change to be solely responsible for an observed biofilm defect. Despite this observation, we did find that a small number of nutrients were responsible for the largest (albeit modest) change in expression for many genes. For example, we identified several groups of genes whose expression changed in the same nutrient conditions (i.e., m-tartaric acid, D-ribose, glycogen), possibly suggesting co-regulation. Given this finding, we speculate that some genes in the c-di-GMP network may be regulated as “suites”, and coordinated 2- to 5-fold changes to a single nutrient across 8 to 12 genes may be an effective method of governing the network by controlling which proteins are present under that given environmental condition. Why would these nutrients in particular regulate suites of genes? Interestingly, m-tartaric acid is produced and metabolized by fluorescent pseudomonads (45) and plays a critical role in solubilizing inorganic phosphate (46); phosphate is a key regulator of biofilm formation by this microbe (47). The physiological relevance of D-ribose and glycogen is less clear. Finally, given that many of the members of the c-di-GMP network, including those encoded by these suites of genes, also have putative ligand binding domains, it is quite possible that more that one mechanism regulates the function and/or specificity of these network members.

Physical interaction was also found to play a role in the network. In comparison to our transcriptional results, we found many discrete examples where two genes, when mutated and shared an altered biofilm phenotype on the same nutrient source also physically interact with one another. Of the 85 pairs of interacting proteins analyzed only 6 shared no biofilm phenotypes, and more than a quarter (24) of the pairs shared a biofilm phenotype in 25 of more nutrient conditions. The remaining pairs shared phenotypes between 1 and 25 nutrient conditions. We interpret the relatively common frequency of interacting pairs sharing biofilm phenotypes to indicate that physical interaction occurs among network members that can exert their influence in similar environmental conditions. Such a combination of common nutrient inputs among interacting proteins provides one of the strongest findings supporting the concept of multimodal regulation. In the case of those interaction pairs which show little or no overlap in their mutant’s biofilm phenotype, it possible that we have not yet identified the correct conditions for which the interaction pair is relevant. Alternatively, some of these enzymes and effectors may interact but are alternatively activated in differing environments.

The potential for physical interaction, especially among the DGCs and the LapD receptor is potentially informative regarding signaling specificity. We reported previously that GcbC and LapD interact (14), and a recent report by Cooley, O’Donnell and Sondermann (48) proposed that LapD forms a dimer-of-dimers “basket”, with space for a DGC to nestle within the basket – it is possible, for example, that all DGCs that interact with LapD have the potential to form this signaling complex. Furthermore, given that many DGCs have associated ligand-binding domains and show relatively few examples of ligand-mediated transcriptional control, we propose that interaction of DGCs with the LapD receptor, coupled with ligand-mediated control of DGC activity, could be a general mechanism of signaling specificity. Finally, the propensity of members of the network to interact indicates the possibility that “local” signaling is a common aspect of the network, and any models describing how the network functions cannot ignore this feature (49).

We also identified a number of putative DGCs that worked ‘against type’, whereby their absence actually resulted in an increase in biofilm formation in a number of conditions at varying frequencies. Previous work has shown that the DGC GcbA in *P. aeruginosa* is partly responsible for modulating a PDE linked to dispersal of biofilm (50), providing some precedence that DGCs do not necessarily have to promote biofilm formation. However, we note here that two DGCs (i.e., Pfl01_2049 and Pfl01_2297) that interact with LapD demonstrated this analogous phenotype under some conditions, leading to the possibility that some DGCs may exert their impact by binding and blocking effector proteins. Additionally, strains carrying mutations in some DGCs were found to either enhance or reduce biofilm formation depending on the nutrient tested, suggesting the mechanism of their impact on biofilm formation of some of these enzymes may be different from what was reported for the *P. aeruginosa* GcbA enzyme. It is not inconceivable that physical interaction may play a role in this phenomenon as well, with DGCs blocking effectors until the correct nutrient/ligand activates the DGC, whereupon catalytic activity is initiated or structural rearrangements cause the enzyme to change its interaction profile.

While we have provided insight into the how a large c-di-GMP network is organized and regulated, we note that there are also several hurdles to interpreting the results presented here. One alternative explanation for the high percentage of c-di-GMP related proteins impacting biofilm is that they are in fact responsible for different cellular process that indirectly have an impact on biofilm formation. It is also likely that the nutrients tested here do not represent the complete collection of environmental cues that these enzymes and effectors respond to in terms of catalytic activation and transcriptional regulation. Indeed, the largest transcriptional change observed was for *rapA*, the PDE up-regulated in a low phosphate condition, suggesting that inputs other than organic compounds can impact the network. Further, we note that several transcripts measured in our Nanostring experiments showed very low expression under most conditions. Still, we find it compelling that a small subset of nutrients produced both the highest and lowest transcriptional changes of a small number of genes, indicating that genes may be regulated as suites in addition to being regulated individually by highly specific input signals. Finally, we note that the bacterial two-hybrid only presents evidence of which proteins interact with each other (or not) in a heterologous host under one growth condition (i.e., LB). That is, it is conceivable that the ligand bound state of a DGC or PDE may not only activate/deactivate enzyme activity, but it may also cause structural changes making the protein competent or incompetent for interaction. Additionally, there may be further interactions we were not able to observe in this system, either because the protein is unstable/unable to function outside of its native host, or alternatively because the interaction was not observable under our assay conditions.

Taken together, our results show that this large c-di-GMP network is a dynamic system capable of responding in specific ways to a variety of inputs with individual members able to take on different functions under differing circumstances. Importantly, we identified numerous instances where combined modes of regulation are observed (nutrients inputs/protein-protein interactions, transcription control/protein-protein interactions, nutrient inputs/transcriptional control). Such multimodal regulation would allow the integration of multiple inputs to fine-tune the output(s) of c-di-GMP-regulated processes, likely enhancing specificity of response by this network. Future studies will be required to examine the detailed mechanisms of ligand recognition, physical interaction among member proteins, and transcriptional regulation of genes in the network.

## Materials and Methods

### Strains and Media

*P. fluorescens* Pfl01, *E. coli* S17 and *E. coli* BTH101 are used throughout this study (51, 52). *P. fluorescens* and *E. coli* BTH101 were grown at 30°C on 1.5% agar LB plates unless otherwise indicated. *E. coli* S17 was grown at 37°C. Media used in this study includes K10T-1 as reported previously (52), and base minimal medium (BMM). BMM is composed of 50 mM Tris HCL (pH 7.4), 7.56 mM (NH_4_)_2_SO_4_, 0.15% glycerol, 1 mM K_2_HP0_4_, and 0.6 mM MgSO_4_. *P. fluorescens* strains harboring the pMQ72 expression plasmid were grown overnight with 10 μg/ml gentamicin. Expression was induced during experimentation with 0.2% arabinose. In-frame deletions were constructed as previously reported (53). Ligation cloning of plasmids was conducted using standard laboratory techniques. All strains and plasmids used in this study are listed in **Table S1**.

### Biofilm Assay

*P. fluorescens* strains were struck out on LB plates overnight. In the case of mutants coming from a previously generated mutant library, plates were supplemented with 30 μg/ml tetracycline (30). Single colonies were picked the following day to grow overnight in liquid LB medium, and supplemented with15 μg/ml tetracycline when appropriate (see **Table S1**). Cells densities were measured and normalized to OD_600_ = 0.716. Cells were mixed 1.5:100 with the BMM. 125 μl of the inoculated BMM was pipetted up and down in the Biolog PM1 and PM2A nutrient plates; 100 μl of this mixture was then transferred from the Biolog plates to a standard 96-well polystyrene plate (Costar) to perform the biofilm assays. Plates were covered and placed in a humidified micro-chamber at 30°C for six hours, at which time the liquid in the wells was discarded and the wells were stained with 125 μl of 0.1% crystal violet (CV) at room temperature for ∼20 minutes, then rinsed 2-3 times with water. Wells were allowed to dry overnight, and were de-stained the following day using 150 μl of a solution of water, methanol, and acetic acid (45:45:10 ratio by volume) for 20 minutes at room temperature. 125 μl of the solubilized CV solution was transferred to a flat-bottom 96 well plate and the OD was recorded at 550 nm.

### Analysis of Biofilm Data

For all analysis of experiments assaying biofilm formation using nutrients from the Biolog plates, OD_550_ values were adjusted in the following ways: First, a baseline staining was established for each condition by growing a strain lacking the genes *gcbA, gcbB, gcbC, and wspR* (referred to as Δ4DGC) and subtracting each of these values from the corresponding values in each condition from all other experiments. Second, the variability of WT values for each conditions were assessed and conditions where WT exhibited a coefficient of variance of greater than 0.35 were excluded from further analysis to reduce errors due to noisy conditions. Third, one batch of data (Batch #11, **Table S2**) was higher than all others so that was batch-normalized by multiplying all data from that day by the ratio between the WT median on that day and the median WT value across all other days.

For all analyses except that for **Figure S2**, mutant values were compared to the mean WT value for each condition and the difference between these values was expressed in terms of the number of standard deviations of WT data in that condition. The portion of values in a normal distribution that would be expected to be this many or more standard deviations from the mean was expressed as a p value. All p values were adjusted using a Benjamini-Hochberg adjustment.

An example calculation of the P value is as follows: WT in carbon source X has mean =0.3 and standard deviation = 0.1. Mutant A has a value of 0.496. Difference between WT mean and value for mutant A = 0.496 - 0.3 = 0.196. The number of WT standard deviations this value differs from WT mean = 0.196 / 0.1 = 1.96, and the portion of normally distributed population one would expect to differ from the mean by 1.96 SDs or more = 0.05. Therefore the unadjusted p value for Mutant A in condition X is 0.05.

For **Figure S2,** OD_550_ values for *P. fluorescens* Pf01 and each mutant in each condition were expressed as a ratio compared to the median OD_550_ value of that mutant across all conditions. This approach serves as a representation of the response to that condition relative to an approximation of the baseline biofilm formation. A second ratio was then calculated by comparing the ratio in each condition to the median of WT ratios in that condition. Using the standard deviation of WT ratios for each condition as described above, we assigned a p value to each value.

### Transcriptional Expression

11 replicates of 5 μl from overnight LB liquid cultures of the wild type and mutant strains were spotted onto 1.5% agar plates containing either K10T-1 medium, or the BMM alone or containing 2 mM of the desired nutrient sources. Exceptions were made for nutrients that have previously been published as having biofilm phenotypes including arginine (0.4%), citric acid (0.4%), and L-arabinose (0.2%), or where exact measurements were infeasible (dextrin; soluble fraction – 0.2%). Cells were grown at 30°C for six hours, scraped from the plate and resuspended in TE buffer, and processed using the RNAeasy extraction kit (Qiagen). 75 ng of each RNA sample was added to the Nanostring nCounter kit, and the protocol provided by the manufacture was followed without modification (Nanostring).

### Transcriptional Analysis

Raw reads from the Nanostring system were normalized as a fraction on the total number of reads measured for targeted genes, expressed as reads per 1000 transcripts as described previously (34). A K10T-1 sample and a base medium sample was processed on each cartridge, yielding six biological replicates. The six replicates of the BMM were used to test significance in the difference between BMM and BMM plus other compounds. This analysis was conducted in the same way as for the biofilm data analysis described above. Briefly, each reading in the BMM plus a compound was compared to the mean value in base medium alone. The standard deviation of that gene in the base medium was then used to estimate the portion of values one would expect to find to be as different from the mean as the value being tested or more extreme. This percentage was taken to be the p value. All p values were adjusted using the Benjamini-Hochberg method. Of the 62 genes analyzed, 48 showed less than 2-fold variation from the lowest to highest count between six replicate experiments when grown on K10T-1 (**Table S11**), and another 12 showed variation between 2 and 3-fold. Two genes showed greater than 3-fold change; these genes were Pfl01_1252 and *gcbC*; both of these genes were found to have low counts across all experiments (**Table S4, S11**).

### Bacterial Two-Hybrid Studies and Analysis

BTH101 cells were electroporated with 50 ng of the puT18 and pKNT25 plasmids containing the construct to be tested. Cells were recovered for 1 hr in 1ml LB at 37°C. 50 μl was then spread on selective agar containing 50 μg/ml carbenicillin, 50 μg/ml kanamycin, 40 μg/ml 5-bromo-4-chloro-3-indolyl-β-D-galactopyranoside (X-gal), and 0.5 mM Isopropyl β -D-1-thiogalactopyranoside (IPTG). Plates were incubated at 30°C for 20 hours, and colonies were examined for a blue color. A leucine zipper protein served as a positive control, and the empty plasmids acted as the negative control. Each gene of interest was fused into both the plasmids, and a positive result in either orientation was recorded as an interaction.

## Acknowledgements

The authors would like to thank Colleen Harty and T. Jarrod Smith for their contributions in developing use of the Biolog plates, James Rudd for consulting on hierarchical clustering methods in R, and Tom Hampton for advice on the statistical analysis of the data.

## Author Contributions

Project was conceptualized by GAO, KMD, and CSG. Data curation, validation, and visualization was conducted by KMD and AJC. Formal Analysis was conducted by KMD, AJC, JNT, and GD. Funding was acquired by GOA, CSG, and DAH. Investigation was conducted by KMD, AJC, and TJG. Methodology was developed by KMD. Project was administered, supervised, and resources provided by GAO. Software was contributed by AJC and JNT. Writing the original manuscript was conducted by KMD, and review and editing was conducted by KMD, AJC, GAO, and CSG.

## Supplemental Figure Legends

**Figure S1. Biofilm phenotypes vary with nutrients.** Shown is a histogram of the quantitified biofilm for the WT as a function of the nutrient added to BMM. The histogram is arranged in order from lowest biofilm formed to highest biofilm formed. The red bar indicated the biofilm formed in BMM.

**Figure S2. Heat map of biofilm phenotypes for each mutant with each nutrient.** In this heatmap, red indicates a higher ratio than WT and blue is lower ratio than WT. That is, cells that are red indicate conditions where a mutant biofilm either increased more or decreased less than WT in a condition relative to the median biofilm formed by that mutant. Black indicates values not significantly different from WT.

**Figure S3. Heat map of gene expression for each gene in the network on 50 different nutrients.** Expression of genes when grown on the indicated nutrient-supplemneted BBM in liquid medium. Only values which are significantly different from expression on BMM are colored (see Materials and Methods for details of statistical tests). Red cells indicate conditions where genes were expressed at significantly higher than in BMM, while blue cells indicate conditions where genes were expressed at significantly lower levels compared to BMM.

**Figure S4. Summary of expression analysis. A.** Shown is the log_2_ expression ratio for each gene in the network across the 50 nutrients tested. The ratio was calculated with dividing the number of mRNA transcripts on BMM plus the indicated nutrient by the number of mRNA transcripts in the BMM. A positive value indicates an increase in expression in response to the nutrient compared to BMM, while a negative value indicates a decrease in expression compared to BMM. All ratios were transformed to their log_2_ values then plotted. Each colored dot indicated the ratio of a different carbon source, with the key to the colors shown on the right. **B.** Log_2_ transformed ratio of expression of genes when cells were grown on a solid medium divided by their expression in the same medium in broth. The carbon source added to the liquid or solid BMM is indicated. * p<0.05 Wilcoxon signed rank test.

**Figure S5. Comparing characteristics of c-di-GMP gene expression between organisms.** *P. fluorescens* Pf01 Nanostring expression data were acquired as described in the Materials and Methods. *P. aeruginosa* PAO1 expression data were retrieved from the NCBI database of published microarray analyses using the tools developed by Tan *et al*. (42). The *P. aeruginosa* PAO1 compendium data were normalized by assigning the highest value in each experiment the value 1, and the lowest value the value 0. All other values in the experiment were scaled linearly between those values using the following formula: for each value x: z_i_ = (x_i_ – min(x) / (max(x) – min(x)) where z_i_ is the normalized value of x_i_. **A-D**. Genes were grouped according by predicted proteins domains into the groups GGDEF, EAL, Dual, HD-GYP, and PilZ for c-di-GMP-related genes. Genes not encoding one of those domains was termed “non-cdg”. In addition, genes in the Pho regulon (42) were included as an example of the characteristics of genes under transcriptional regulation. **A-B.** The median normalized expression of each gene was calculated and the distribution of those values is plotted as a boxplot with genes grouped according to the characterization of the domains they encode. Panel **A** shows data from *P fluorescens* and panel **B** from *P. aeruginosa* PAO1. **C-D.** The coefficient of variance of the normalized expression of each gene was calculated by dividing the standard deviation by the mean, and the distribution of those values is plotted as a boxplot with genes grouped according to the characterization of the domain-containing proteins they encode. Panel **C** shows data from *P fluorescens* and panel **D** from *P. aeruginosa* PAO1. **E-F.** Spearman correlation values for each pair of c-di-GMP-related gene in each organism were calculated and plotted using the R package “corrplot”. Significance of these correlations was calculated using the “corrtest” function in this package and p values were adjusted using a Bonferroni correction. Correlations with adjusted p values below 0.05 were not plotted. Panel **E** shows data from *P fluorescens* and panel **E** from *P. aeruginosa* PAO1.

**Figure S6. Schematic of protein-protein interaction sub-networks. A.** Shown is the key for each domain type shown in panels B-E. **B-E**. Small interaction networks showing proteins that interact with a large number of other proteins. Such “hub” proteins are Pfl0_0192 (panel B), Pfl0_2709 (panel C), Pfl0_2525 (panel D) and Pfl0_4068 (panel E). A summary of all such interactions is presented in **Table S10** and **Figure 4**.

## Literature Cited

1. Kulasakara H, Lee V, Brencic A, Liberati N, Urbach J, Miyata S, Lee DG, Neely AN, Hyodo M, Hayakawa Y, Ausubel FM, Lory S. 2006. Analysis of Pseudomonas aeruginosa diguanylate cyclases and phosphodiesterases reveals a role for bis-(3’-5’)-cyclic-GMP in virulence. Proc Natl Acad Sci U S A 103:2839–2844.

2. Tischler AD, Camilli A. 2004. Cyclic diguanylate (c-di-GMP) regulates Vibrio cholerae biofilm formation. Mol Microbiol 53:857–869.

3. Paul K, Nieto V, Carlquist WC, Blair DF, Harshey RM. 2010. The c-di-GMP binding protein YcgR controls flagellar motor direction and speed to affect chemotaxis by a "backstop brake" mechanism. Molecular Cell 38:128–139.

4. Abel S, Bucher T, Nicollier M, Hug I, Kaever V, Abel Zur Wiesch P, Jenal U. 2013. Bi-modal distribution of the second messenger c-di-GMP controls cell fate and asymmetry during the caulobacter cell cycle. PLoS Genet 9:e1003744.

5. Pultz IS, Christen M, Kulasekara HD, Kennard A, Kulasekara B, Miller SI. 2012. The response threshold of Salmonella PilZ domain proteins is determined by their binding affinities for c-di-GMP. Mol Microbiol 86:1424–1440.

6. Christen M, Kulasekara HD, Christen B, Kulasekara BR, Hoffman LR, Miller SI. 2010. Asymmetrical distribution of the second messenger c-di-GMP upon bacterial cell division. Science 328:1295–1297.

7. Sudarsan N, Lee ER, Weinberg Z, Moy RH, Kim JN, Link KH, Breaker RR. 2008. Riboswitches in eubacteria sense the second messenger cyclic di-GMP. Science 321:411–413.

8. Hickman JW, Harwood CS. 2008. Identification of FleQ from Pseudomonas aeruginosa as a c-di-GMP-responsive transcription factor. Mol Microbiol 69:376–389.

9. Perez-Mendoza D, Coulthurst SJ, Sanjuan J, Salmond GP. 2011. N-Acetylglucosamine- dependent biofilm formation in Pectobacterium atrosepticum is cryptic and activated by elevated c-di-GMP levels. Microbiology 157:3340–3348.

10. Hickman JW, Tifrea DF, Harwood CS. 2005. A chemosensory system that regulates biofilm formation through modulation of cyclic diguanylate levels. Proc Natl Acad Sci U S A 102:14422–14427.

11. Krasteva PV, Fong JC, Shikuma NJ, Beyhan S, Navarro MV, Yildiz FH, Sondermann H. 2010. Vibrio cholerae VpsT regulates matrix production and motility by directly sensing cyclic di-GMP. Science 327:866–868.

12. Bobrov AG, Kirillina O, Forman S, Mack D, Perry RD. 2008. Insights into Yersinia pestis biofilm development: topology and co-interaction of Hms inner membrane proteins involved in exopolysaccharide production. Environ Microbiol 10:1419–1432.

13. Lindenberg S, Klauck G, Pesavento C, Klauck E, Hengge R. 2013. The EAL domain protein YciR acts as a trigger enzyme in a c-di-GMP signalling cascade in E. coli biofilm control. EMBO J 32:2001–2014.

14. Dahlstrom KM, Giglio KM, Collins AJ, Sondermann H, O’Toole GA. 2015. Contribution of physical interactions to signaling specificity between a diguanylate cyclase and its effector. MBio 6: e01978–01915.

15. Monds RD, Newell PD, Gross RH, O’Toole GA. 2007. Phosphate-dependent modulation of c-di-GMP levels regulates Pseudomonas fluorescens Pf0-1 biofilm formation by controlling secretion of the adhesin LapA. Mol Microbiol 63:656–679.

16. Lee SH, Angelichio MJ, Mekalanos JJ, Camilli A. 1998. Nucleotide sequence and spatiotemporal expression of the Vibrio cholerae vieSAB genes during infection. J Bacteriol 180:2298–2305.

17. Lim B, Beyhan S, Meir J, Yildiz FH. 2006. Cyclic-diGMP signal transduction systems in Vibrio cholerae: modulation of rugosity and biofilm formation. Mol Microbiol 60: 331–348.

18. Yildiz FH, Liu XS, Heydorn A, Schoolnik GK. 2004. Molecular analysis of rugosity in a Vibrio cholerae O1 El Tor phase variant. Mol Microbiol 53:497–515.

19. Tuckerman JR, Gonzalez G, Sousa EHS, Wan XH, Saito JA, Alam M, Gilles-Gonzalez MA. 2009. An oxygen-sensing diguanylate cyclase and phosphodiesterase couple for c- di-GMP control. Biochemistry 48:9764–9774.

20. Zahringer F, Lacanna E, Jenal U, Schirmer T, Boehm A. 2013. Structure and signaling mechanism of a zinc-sensory diguanylate cyclase. Structure 21:1149–1157.

21. Dahlstrom KM, O’Toole GA. 2017. A symphony of cyclases: Specificity in diguanylate cyclase signaling. Annu Rev Microbiol doi:10.1146/annurev-micro-090816–093325.

22. Boyd CD, O’Toole GA. 2012. Second messenger regulation of biofilm formation: breakthroughs in understanding c-di-GMP effector systems. Annu Rev Cell Dev Biol 28:439–462.

23. Newell PD, Monds RD, O’Toole GA. 2009. LapD is a bis-(3’,5’)-cyclic dimeric GMP-binding protein that regulates surface attachment by Pseudomonas fluorescens Pf0-1. Proc Natl Acad Sci U S A 106:3461–3466.

24. Newell PD, Boyd CD, Sondermann H, O’Toole GA. 2011. A c-di-GMP effector system controls cell adhesion by inside-out signaling and surface protein cleavage. PLoS Biol 9:e1000587.

25. Liu N, Pak T, Boon EM. 2010. Characterization of a diguanylate cyclase from Shewanella woodyi with cyclase and phosphodiesterase activities. Mol Biosyst 6:1561–1564.

26. Navarro MV, De N, Bae N, Wang Q, Sondermann H. 2009. Structural analysis of the GGDEF-EAL domain-containing c-di-GMP receptor FimX. Structure 17:1104–1116.

27. Navarro MV, Newell PD, Krasteva PV, Chatterjee D, Madden DR, O’Toole GA, Sondermann H. 2011. Structural basis for c-di-GMP-mediated inside-out signaling controlling periplasmic proteolysis. PLoS Biol 9:e1000588.

28. Ryjenkov DA, Simm R, Romling U, Gomelsky M. 2006. The PilZ domain is a receptor for the second messenger c-di-GMP: the PilZ domain protein YcgR controls motility in enterobacteria. J Biol Chem 281:30310–30314.

29. Pratt JT, Tamayo R, Tischler AD, Camilli A. 2007. PilZ domain proteins bind cyclic diguanylate and regulate diverse processes in Vibrio cholerae. J Biol Chem 282:12860–12870.

30. Newell PD, Yoshioka S, Hvorecny KL, Monds RD, O’Toole GA. 2011. Systematic analysis of diguanylate cyclases that promote biofilm formation by Pseudomonas fluorescens Pf0-1. J Bacteriol 193:4685–4698.

31. Townsley L, Yildiz FH. 2015. Temperature affects c-di-GMP signalling and biofilm formation in Vibrio cholerae. Environ Microbiol doi:10.1111/1462-2920.12799.

32. Ha DG, Richman ME, O’Toole GA. 2014. Deletion mutant library for investigation of functional outputs of cyclic diguanylate metabolism in Pseudomonas aeruginosa PA14. Appl Environ Microbiol 80:3384–3393.

33. Spurbeck RR, Tarrien RJ, Mobley HL. 2012. Enzymatically active and inactive phosphodiesterases and diguanylate cyclases are involved in regulation of Motility or sessility in Escherichia coli CFT073. MBio 3.

34. Hollomon JM, Grahl N, Willger SD, Koeppen K, Hogan DA. 2016. Global role of cyclic AMP signaling in pH-dependent responses in Candida albicans. mSphere 1.

35. Luo Y, Zhao K, Baker AE, Kuchma SL, Coggan KA, Wolfgang MC, Wong GC, O’Toole GA. 2015. A hierarchical cascade of second messengers regulates Pseudomonas aeruginosa surface behaviors. MBio 6.

36. Guvener ZT, Harwood CS. 2007. Subcellular location characteristics of the Pseudomonas aeruginosa GGDEF protein, WspR, indicate that it produces cyclic-di-GMP in response to growth on surfaces. Mol Microbiol 66:1459–1473.

37. Kuchma SL, Griffin EF, O’Toole GA. 2012. Minor pilins of the type IV pilus system participate in the negative regulation of swarming motility. J Bacteriol 194:5388–5403.

38. Petrova OE, Cherny KE, Sauer K. 2014. The Pseudomonas aeruginosa diguanylate cyclase GcbA, a homolog of P. fluorescens GcbA, promotes initial attachment to surfaces, but not biofilm formation, via regulation of motility. J Bacteriol 196:2827–2841.

39. Baraquet C, Harwood CS. 2013. Cyclic diguanosine monophosphate represses bacterial flagella synthesis by interacting with the Walker A motif of the enhancer-binding protein FleQ. Proc Natl Acad Sci U S A 110:18478–18483.

40. Lee ER, Baker JL, Weinberg Z, Sudarsan N, Breaker RR. 2010. An allosteric self-splicing ribozyme triggered by a bacterial second messenger. Science 329:845–848.

41. Dahlstrom KM, Giglio KM, Sondermann H, O’Toole GA. 2016. The inhibitory site of a diguanylate cyclase is a necessary element for interaction and signaling with an effector protein. J Bacteriol 198:1595–1603.

42. Tan J, Doing G, Lewis KA, Price CE, Chen KM, Cady KC, Perchuk B, Laub MT, Hogan DA, Greene CS. 2017. Unsupervised extraction of stable expression signatures from public compendia with an ensemble of neural networks. Cell Syst 5:63–71 e66.

43. Baker AE, Diepold A, Kuchma SL, Scott JE, Ha DG, Orazi G, Armitage JP, O’Toole GA. 2016. PilZ domain protein FlgZ mediates cyclic di-GMP-dependent swarming motility control in Pseudomonas aeruginosa. J Bacteriol 198:1837–1846.

44. Andrade MO, Alegria MC, Guzzo CR, Docena C, Rosa MC, Ramos CH, Farah CS.2006. The HD-GYP domain of RpfG mediates a direct linkage between the Rpf quorum-sensing pathway and a subset of diguanylate cyclase proteins in the phytopathogen Xanthomonas axonopodis pv citri. Mol Microbiol 62:537–551.

45. Allen RH, Jakoby WB. 1969. Tartaric acid metabolism. IX. Synthesis with tartrate epoxidase. J Biol Chem 244:2078–2084.

46. Hurlbert RE, Jakoby WB. 1965. Tartaric acid metabolism. I. Subunits of L(+)-tartaric acid dehydrase. J Biol Chem 240:2772–2777.

47. Park KH, Lee CY, Son HJ. 2009. Mechanism of insoluble phosphate solubilization by Pseudomonas fluorescens RAF15 isolated from ginseng rhizosphere and its plant growth-promoting activities. Lett Appl Microbiol 49:222–228.

48. Cooley RB, O’Donnell JP, Sondermann H. 2016. Coincidence detection and bi-directional transmembrane signaling control a bacterial second messenger receptor. Elife 5.

49. Yan J, Deforet M, Boyle KE, Rahman R, Liang R, Okegbe C, Dietrich LEP, Qiu W, Xavier JB. 2017. Bow-tie signaling in c-di-GMP: Machine learning in a simple biochemical network. PLoS Comput Biol 13:e1005677.

50. Petrova OE, Cherny KE, Sauer K. 2015. The diguanylate cyclase GcbA facilitates Pseudomonas aeruginosa biofilm dispersion by activating BdlA. J Bacteriol 197:174–187.

51. Simon R, Priefer U, Puhler A. 1983. A broad host range mobilization system for invivo genetic-engineering -Transposon mutagenesis in gram-negative bacteria. Bio-Technology 1:784–791.

52. Monds RD, Newell PD, Schwartzman JA, O’Toole GA. 2006. Conservation of the Pho regulon in Pseudomonas fluorescens Pf0-1. Appl Environ Microbiol 72:1910–1924.

53. Shanks RM, Caiazza NC, Hinsa SM, Toutain CM, O’Toole GA. 2006. Saccharomyces cerevisiae-based molecular tool kit for manipulation of genes from gram-negative bacteria. Appl Environ Microbiol 72:5027–5036.

